# COVID-19: A need for new rather than repurposed antiviral drugs

**DOI:** 10.1101/2021.03.25.436935

**Authors:** Dory Kovacs, Chris Davis, Paul Cannon, Melanie McFarlane, Stephanie M Rainey, Rute Pinto, Meredith E Stewart, Agnieszka M Szemiel, Aislynn Taggart, Alain Kohl, Fiona Marra, Emma C Thomson, Janet T Scott

## Abstract

**Background:** SARS-CoV-2 infection, the causative agent of COVID-19, has resulted in over 2,500,000 deaths to date^1^. Although vaccines are becoming available, treatment options remain limited. Repurposing of compounds could reduce the time, cost, and risks associated with the development of new drugs and has been the focus of many clinical studies.

Here, we summarise available evidence on 29 FDA-approved compounds, from in vitro results to clinical trials, focussing on remdesivir, galidesivir and favipiravir, and test 29 antiviral compounds’ activity *in vitro*.

**Methods:** A comprehensive search strategy was used to retrieve trials and publications related to antiviral compounds with potential efficacy to treat coronaviruses. These data were used to prioritise testing of a panel of antiviral drugs in vitro against patient isolates of SARS-CoV-2. An in vitro screen was carried out to determine the activity of 29 FDA-approved compounds.

**Results:** 625 clinical trials investigated 16 repurposed antiviral candidate compounds for the treatment of COVID-19. In vitro studies identified ten drug candidates with demonstrable anti-SARS-CoV-2 activity, including favipiravir, remdesivir, and galidesivir. To validate these findings, a drug screen was conducted using two cell lines and wildtype isolates of SARS-CoV-2 isolated from patients in the UK. While eight drugs with anti-SARS-CoV-2 activity were identified in vitro, activity in clinical trials has, as yet failed to demonstrate a strong effect on mortality.

**Conclusions:** So far, no repurposed antiviral has shown a strong effect on mortality in clinical studies. The urgent need for novel antivirals in this pandemic is clear, despite the costs and time associated with their development.

**Research in Context:** *Evidence before this study:* Repurposing of existing compounds for the treatment of COVID-19 has been the focus of many *in vitro* studies and clinical trials, saving time, costs and risks associated with the research and development of new compounds.

*Added value of this study:* We reviewed the literature for 29 FDA-approved compounds with previously reported (or suspected) anti-SARS-CoV-2 activity and found 625 clinical trials that have been undertaken on 16 different drugs. We determined if repurposed antivirals are suitable for clinical trials based on previously published data, and conducted an additional *in vitro* screen using locally circulating strains in the UK (PHE2 and GLA1). We report the difference in IC_50_ from published data using Wuhan1/Wash1 strains with PHE2 and GLA1, including IC_50_ values below 100μM for galidesivir in wild-type virus. Given the limited success of repurposed compounds in the treatment of COVID-19, we comment on the urgent need for new antivirals specifically targeting SARS-CoV-2.

*Implications of all the available evidence:* Our data show that most prospective compounds for repurposing show no anti-SARS-CoV-2 activity, and antiviral activity *in vitro* does not always translate to clinical benefit. So far, no repurposed compound has shown a strong effect on mortality in clinical studies. Drugs, including monoclonal antibody therapies, that have been developed to target SARS-CoV-2 virus itself have shown most promise.

## 1 Introduction

The betacoronavirus SARS-CoV-2 is likely to be a significant public health threat to the global population in the medium to long term. This coronavirus family of RNA viruses are commonly associated with mild to moderate upper respiratory tract infections during the winter months^2,3^. However, the emerging coronaviruses MERS-CoV^4^, SARS-CoV-1^5^ and SARS-CoV-2^6^ have high mortality rates. In addition, long-term morbidity associated with SARS-CoV-2 infection, known as ‘long COVID’^7^ is a significant concern and highlights the need for intervention with antiviral therapies. The novel coronavirus SARS-CoV-2 originated in China in late 2019 and has spread across the globe, resulting in over 2,500,000 recorded deaths to date.

Repurposing of existing compounds for the treatment of COVID-19 has been the focus of many trials and studies, despite a lack of evidence regarding their effectiveness ^8^. Repurposing of compounds is made possible by shared molecular pathways across diseases, which also overcomes bottlenecks in the developmental process by cutting the time, costs, and risks associated with drug development^9-11^. Previously developed antivirals may demonstrate activity against diverse viral families due to the presence of relatively conserved viral proteins such as DNA or RNA polymerases, integrase or viral proteases. Many viruses also require inactivation of host proteins, which could also provide targets for broad acting antivirals. Longstanding compounds have the advantage of pre-existing information regarding their pharmacokinetic and pharmacodynamic profiles and toxicity. However, repurposing drugs and compounds is not without caveats, and must occur in controlled, clinical trial settings. While a drug may be efficacious against the original target, the concentration required to combat another virus may be unachievable^12^ due to toxicity or pharmacodynamic constraints in patients. Furthermore, there is a finite number of antivirals to trial, and emerging evidence is often conflicting. As a result, clinicians often make decisions in the absence of reliable data, based on preliminary reports and poorly conducted trials.

Herein, we summarise evidence on 29 FDA-approved candidate compounds, from in vitro results to clinical trials selected for potential SARS-CoV-2 antiviral activity. These potential antiviral candidates were next tested for activity against circulating SARS-CoV-2 variants in vitro.

## 2 Methods

### 2.1 Literature review

Antiviral drugs and compounds were identified by screening literature on SARS, MERS and the emergent literature on COVID-19, coupled with the experience of the authors (see Table 1). We then undertook a comprehensive search strategy to retrieve trials and publications related to antiviral drugs and compounds potentially effective in the treatment of coronaviruses. The method was adapted from that recommended in the Cochrane Handbook for Systematic Reviews of Interventions^13^ for identifying trials. Searches for the below list of antiviral drugs and compounds were conducted between March 2020 and 28 January 2021 on ClinicalTrials.gov, the WHO International Clinical Trials Registry Platform (ICTRP) and the Cochrane COVID-19 Study Register. Search terms are shown in Supplementary Table 1. In addition, we searched the Chinese Clinical Trial Register and pre-print servers medRxiv, bioRxiv, and chinaRxiv, while relevant results through our academic network were also identified.

**Table 1.** Twenty-nine compounds were identified for reports of anti-SARS-CoV-2 activity. Drugs with no oral formulations are shown in orange, original indications are shown in brackets. HBV=hepatitis B virus, HCV=hepatitis C virus, HIV = human immunodeficiency virus, EVD = Ebola virus disease, RSV= respiratory syncytial virus.

| Nucleoside analogues | Protease inhibitors | Anti-influenza drugs | Others |
| --- | --- | --- | --- |
| Abacavir (HIV) | Atazanavir (HIV) | Amantadine | Emetine |
| Adefovir (HBV) | Danoprevir (HCV) | Baloxavir | Hydroxychloroquine |
| Didanosine (HIV) | Darunavir (HIV) | Favipiravir*** | Sotetsuflavone |
| Emtricitabine (HIV, HBV) | Fosamprenavir (HIV) | Oseltamivir |  |
| Galidesivir (HCV*) | Grazoprevir (HCV) | Umifenovir** |  |
| Lamivudine (HIV, HBV) | Lopinavir (HIV) |  |  |
| Remdesivir (EVD) | Paritaprevir (HCV) |  |  |
| Ribavirin (HCV, RSV) | Ritonavir (HIV) |  |  |
| Sofosbuvir (HCV) | Saquinavir (HIV) |  |  |
| Telbivudine (HBV) |  |  |  |
| Tenofovir (HIV, HBV) |  |  |  |
| Zidovudine (HIV) |  |  |  |

\*Galidesivir was subsequently used for filovirus infections, including Ebola<sup>14</sup>. \*\*Umifenovir is also known as Arbidol. \*\*\*Favipiravir is a nucleoside analogue.

On the Cochrane COVID-19 Study Register, free text terms were used to search for the name of each drug or compound individually; no other search terms or limits were applied. This process was repeated by searching medRxiv and bioRxiv simultaneously through bioRxiv in the title or abstract fields, and for ChinaRxiv across all fields. For ClinicalTrials.gov and the Chinese Clinical Trial Register, each drug or compound was searched in the intervention field. Results were filtered for relevance before being exported and analysed in EndNote X9. Results were also screened to include only those published in English. No date or language limits were applied to the searches.

Weekly ICTRP reports were compiled and screened for trials mentioning COVID-19 or SARS-CoV-2 in their scientific or public titles, and their details were extracted using R version 4·0·0.

### 2.2 In vitro screen of 29 FDA-approved compounds

#### Cell lines

The methods of production and validation of the cell lines used in this study are fully described elsewhere^15^. In brief, VeroE6 (African green monkey kidney) cells obtained from Prof. Michèle Bouloy (Institut Pasteur, France) were induced to overexpress ACE2 using a modified lentiviral expression system (pLV-EF1a-IRES-Neo (Addgene, plasmid 85139)). A549 cells (human lung) expressing the NPro protein of bovine viral diarrhea virus (BVDV) were obtained from Prof. Rick Randall (University of St Andrews) and induced to overexpress ACE2 in the same manner as the VeroE6 cells.

#### Virus isolates and propagation

Two clinical isolates were used in the study, including the PHE2 isolate, provided by Public Health England, and a clinical isolate obtained from the MRC-University of Glasgow Centre for Virus Research, Scotland, UK (GLA1). GLA1 was obtained by inoculating a T-25 flask of VeroE6 cells with a bronchoalveolar lavage clinical sample and incubating the cells until CPE was observed. Both stocks were confirmed using metagenomic sequencing and expanded by inoculating a T-150 cm^2^ of VeroE6 cells and incubating for 48 hours before clarifying the supernatant. A plaque assay was used to determine an accurate titre for each virus and ensure an accurate multiplicity of infection (MOI) for each isolate in subsequent experiments.

#### Endpoint Dilution Assay

An in vitro screen was performed to determine the activity of 29 FDA-approved compounds (see Table 1) on the ability of SARS-CoV-2 to infect and cause cytopathic effect (CPE) on the two cell lines derived from A549 and VeroE6 cells, as described above. A549-NPro-ACE2 and VeroE6-ACE2 cells were plated at a density of 1·25×10^4^ cells/well in a 96-well plate in DMEM supplemented with 10% FCS the day before drug treating the cells. The media was replaced with 50ul DMEM-GlutaMAX (ThermoFisher, cat no 10566016) supplemented with 2% FCS (ThermoFisher, cat no A4766801) containing doubling dilutions of the various compounds at the stated concentrations. The cells were incubated with the compounds for three hours before the addition of 50ul of either SARS-CoV-2 PHE2 or GLA1 isolates at a MOI of 0·05 or 0·5. The concentrations presented in the results equate to the concentration of the compounds after the addition of the virus isolates. 72 hours post infection the cells were fixed with 8% (v/v) formaldehyde in PBS for one hour, washed with PBS and then stained with Coomassie solution (2ml Coomassie Brilliant Blue (ThermoFisher cat no. 20277), 900ml Methanol, 900ml water 200ml Acetic Acid) for 30 minutes. The plates were washed in water to remove the stain and dried overnight before using a Celigo Imaging Cytometer (Nexcelom Bioscience, UK) to measure the staining intensity. Percentage cell survival was determined by comparing the intensity of the staining to an uninfected well.

#### Statistical packages

All graphs were produced using Prism (GraphPad, version 8) with the IC_50_ values determined by a nonlinear regression using a sigmoidal 4PL equation.

## 3 Results

### 3.1 Clinical trials with repurposed antivirals

The WHO’s ITCRP database was reviewed for clinical trials examining the repurposing of 29 compounds for the treatment of COVID-19. Between March 2020 and January 2021, 655 such trials were registered, investigating sixteen drugs in mild to severe COVID-19 disease. For trials with registered details, median target size was 135 patients. Many trials were multi-national with multiple experimental arms. Iran had the highest number of clinical trials registered (n=145), followed by the US (n=72) (Figure 1). In Europe, Spain had the highest number of trials registered (n=40). 68 studies investigating drug repurposing in China, were initiated but concluded early due to a lack of patients. The number of trials remained low in Africa; however, one multicentre trial (PACTR202006537901307) investigated several antiviral compounds across 13 nations on the continent.

**Figure 1.**
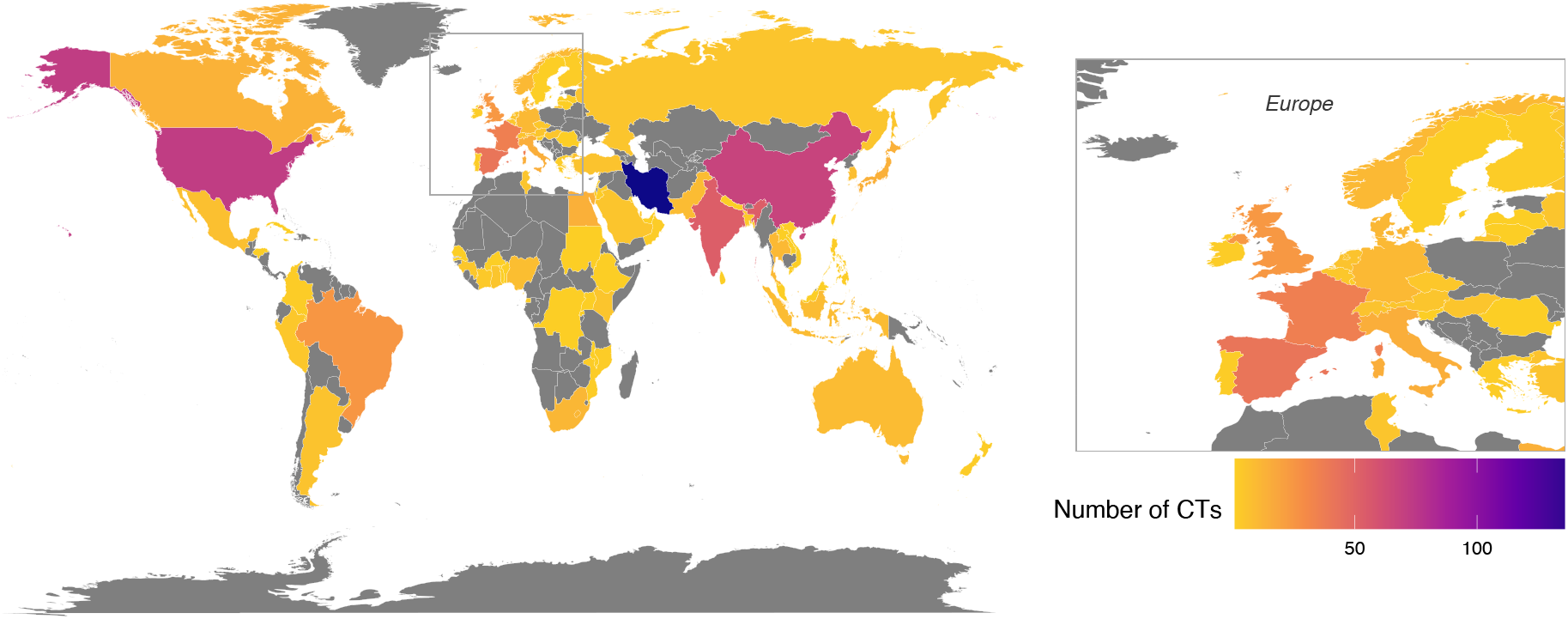
Number of clinical trials looking at repurposing of compounds for the treatment of COVID-19 per country. Only trials that specified countries are shown (n=614). Source: WHO ITCRP database, March 2020 - January 2021.

The most commonly repurposed inhibitor was HCQ (n=461, see Figure 2), a known anti-malarial and blocker of virus entry, which can be cheaply produced and dispensed around the globe. HCQ, lopinavir and ritonavir (the latter two are co-formulated) were investigated early in the pandemic (up to July 2020) but have now been removed from trials in hospitalised patients following data from the WHO’s Solidarity trial in which they were shown to have no effect on mortality^16-18^.

**Figure 2.**
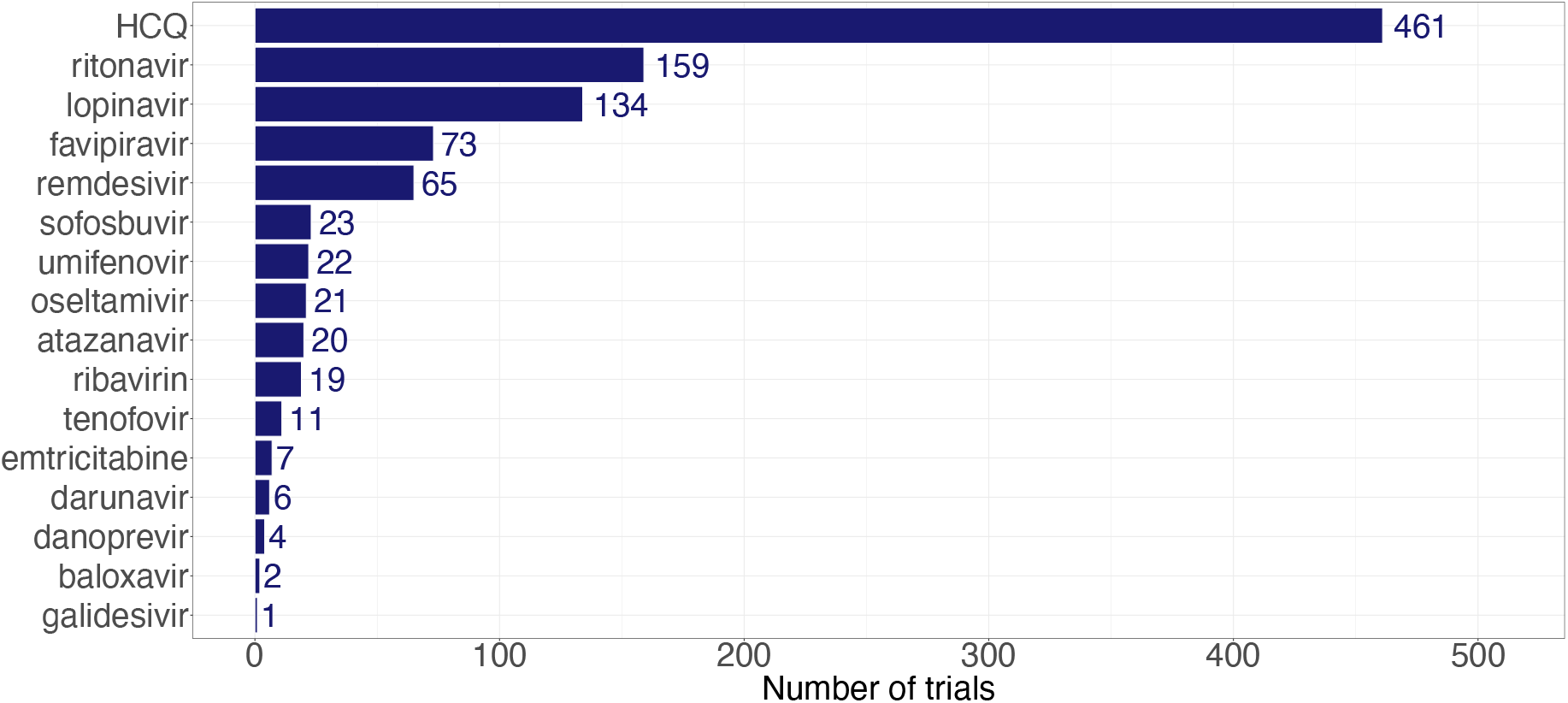
Number of trials looking at the repurposing of sixteen compounds as of January 2021. HCQ = hydroxychloroquine; (Although there were 655 unique trials registered, the total here adds up to 1028 as many trials investigate multiple compounds through multiple experimental arms.)

Data on the antiviral drug remdesivir, previously tested in patients with Ebola virus disease were initially encouraging, with evidence that time to recovery was reduced in hospitalised patients^19^. However, these results were not replicated in the Solidarity trial^17,20^, which also found no effect of HCQ, lopinavir/ritonavir or interferon on overall mortality, initiation of ventilation or duration of hospital stay in hospitalized patients. These conflicting results may reflect the need to initiate antiviral treatment early before the onset of pathology driven by the immune response.

Favipiravir was the subject of 73 trials. Two trials have reported a faster time to viral clearance in favipiravir treated patients compared to controls (^21^, FUJIFILM Toyama Chemical press release). Two trials are ongoing in the UK: Getafix^22^ and Pioneer^23^.

The first clinical trial assessing galidesivir in COVID-19 commenced in April 2020^14^ (NCT03891420) and is expected to be completed by May 2021.

### 3.2 Repurposed compounds and anti-SARS-CoV-2 activity in vitro

Of 29 compounds (Table 1), ten drug candidates have repeatedly been shown to have anti-SARS-CoV-2 activity in vitro. These compounds and their EC_50_ and IC_50_ values are shown in Figure 3 (and Supp Table 1). In addition, Yamamoto et al^24^ reported that saquinavir and atazanavir inhibited SARS-CoV-2 at 8·83μM and 9·36μM, but the Cmax/EC_50_ and _Cmax/_C_trough_ ratio was not >=1, and therefore it was considered unlikely that these drugs would be useful clinically (saquinavir C_max_/EC_50_ = 0·73, C_trough_/EC_50_ = 0·07, atazanavir C_max_/EC_50_ = 0·93, C_trough_/EC_50_ = 0·19). Remdesivir has been shown to have high antiviral activity in vitro, with consistently low EC_50_ values in VeroE6^25-28^, Caco-2^29-31^, Calu-3^32,33^ and Huh7.5^34^ cells. HCQ is active against SARS-CoV-2 in VeroE6, Huh7.5 and Calu-3 cells^34-37^ and is the most commonly investigated compound in clinical trials (Figure 2 and 3). Lopinavir, frequently used in combination with ritonavir, has been shown to inhibit SARS-CoV-1^38,39^ and has some activity against SARS-CoV-2^24,25,30,32,40^ in Caco-2, Calu-3 and VeroE6 cells. Several studies reported no antiviral effect associated with favipiravir under 100μM in VeroE6 cells^25,40,41^, Wang et al^26^ reported a single EC_50_ of 61·88 μM indicating an inhibitory effect similar to that for Ebola virus. Galidesivir has been shown to have antiviral activity against SARS-CoV-1 and MERS-CoV in HeLa cells^42^ and against culture-adapted SARS-CoV-2^43^. Ribavirin and sofosbuvir have EC_50_ values under 10μM in Huh7.5 and Calu-3 cells^44^. Emetine, used for the treatment of amoebiasis, has been shown to inhibit MERS-CoV and SARS-CoV-2 in vitro^25,45^. The hepatitis C virus proteases grazoprevir and danoprevir have also shown anti-SARS-CoV-2 activity with EC_50_s below 100 μM^46^, while danoprevir is also investigated in four clinical trials. Information relevant for clinical use of these compounds are summarised in Supplementary Table 3.

**Figure 3.**
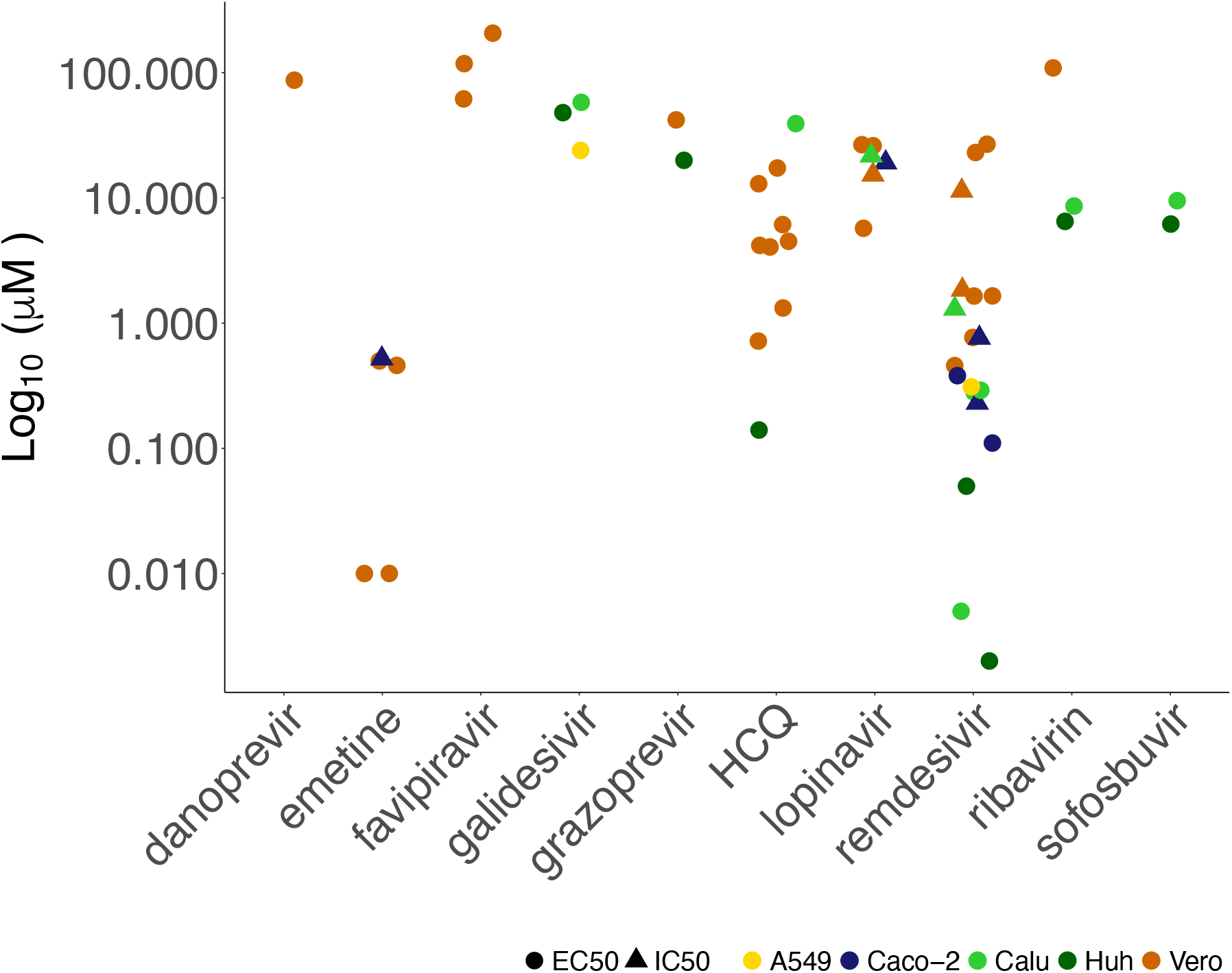
EC_50_ and IC_50_(log_10_) values of selected compounds reported in previous publications. Note that MOIs varied from 0.001 to 1; see Supplementary Table 2.

### 3.3 Galidesivir, remdesivir and hydroxychloroquine demonstrate high antiviral activity against SARS-CoV-2 in vitro

The antiviral activity of 29 FDA-approved compounds was tested in vitro, using VeroE6-ACE2 and A549-NPro-ACE2 cells. Of these, eight showed activity against SARS-CoV-2 PHE2 in VeroE6-ACE2 cells to varying degrees (Supp Figure 4).

Galidesivir, remdesivir and HCQ were able to increase the survival of VeroE6-ACE2 when infected with the GLA1 and the PHE2 isolates (Supp figure 2, Supp figure 3). Lopinavir, ribavirin and favipiravir reduced cell death in both cell lines to a small degree at the highest concentrations, but this coincided with a cytotoxicity, with the latter two only showing inhibition at the highest concentrations of the drugs tested against an MOI of 0·05, with little effect on the virus at an MOI of 0·5. The same effect was seen with emetine and umifenovir with both increased cell survival but at concentrations that also lead to greater cell cytotoxicity than seen in lopinavir, ribavirin and favipiravir.

The compounds were then tested on A549-NPro-ACE2 cells, which constitute a more physiologically relevant human cell line that is susceptible to SARS-CoV-2 virus infection. The results were very similar to those seen in VeroE6-ACE2 cells apart from favipiravir, which had no apparent effect (Figures 4). Again, ribavirin showed an ability to increase cell survival against the PHE2 virus but not the GLA1 isolate (Supp figure 1 and Figure 4, respectively).

**Figure 4.**
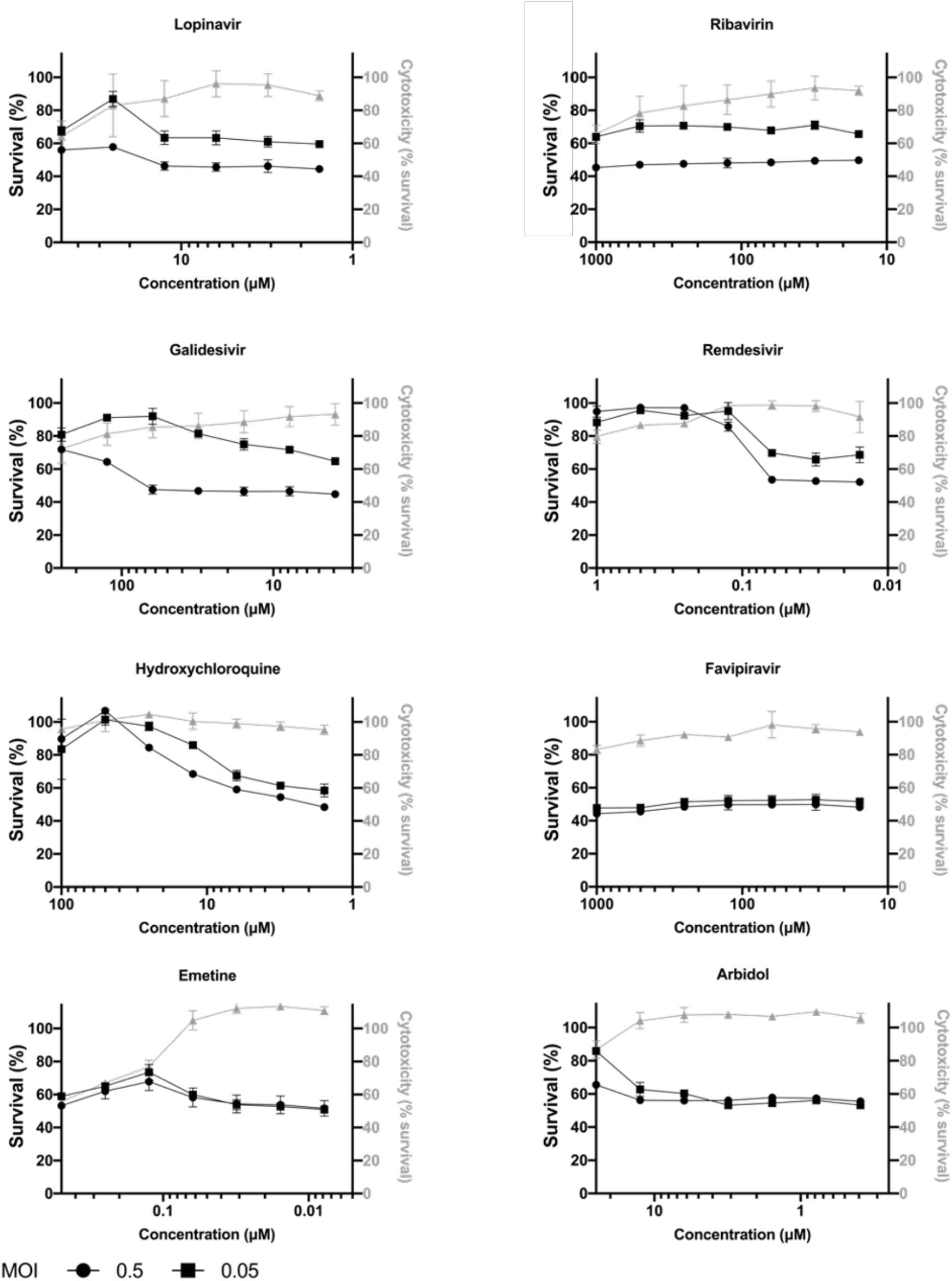
Assessment of SARS-CoV-2 inhibition by active FDA approved antiviral compounds. A549 cells, overexpressing ACE2 and BVDV Npro protein, were treated for three hours with varying concentrations of antiviral compounds before the addition of SARS-CoV-2 GLA1 isolate at an MOI of 0·05 and 0·5. The cells were fixed with 8% formaldehyde 72 hours post-infection and stained with Coomassie blue. Percentage cell survival was determined by comparing the intensity of the staining to uninfected and untreated wells. The data shown are representative figures of 3 independent repeats. The mean percentage cell survival of the DMSO control infected with an MOI of 0·05 and 0·5 was 50% (SD 0·502) and 43·45% (SD 0·642), respectively. The cytotoxicity of the compounds is depicted by the grey line and were determined by comparing the intensity of staining to untreated wells

IC_50_ values were determined for the three most active compounds against the PHE2 and GLA1 viral isolates on t VeroE6-ACE2 and A549-NPro-ACE2 cells (Table 2). At an MOI of 0·5, galidesivir had an IC_50_ of between 60·5 μM to 106·5 μM, while the IC_50_ of HCQ ranged from 12·53 μM to 92·8 μM, depending on the viral isolate and cell line. Remdesivir demonstrated an IC_50_ of 3·17 μM for the PHE2 and 2·62 μM for the GLA1 isolate on VeroE6-ACE2 cells. On A549-NPro-ACE2 cells the activity was even greater at 0·25 μM and 0·11 μM for the PHE2 and GLA1 cells, respectively. Overall, these results demonstrate that remdesivir and HCQ and, for the first time, galidesivir are able to significantly increase cell survival, while other compounds only show a minor activity against SARS-CoV-2 wildtype isolates, in vitro.

**Table 2.** IC_50_ values (mM) of the three most active compounds against SARS-CoV-2 (PHE 2 and GLA1 isolates). HCQ = hydroxychloroquine; MOI = multiplicity of infection. R^2^, the correlation coefficient, indicates the fit between the datapoints and fitted line (where R^2^=1 is a perfect fit).

|  | VeroE6-ACE2 |  |  |  |  | A549-NPro-ACE2 |  |  |  |
| --- | --- | --- | --- | --- | --- | --- | --- | --- | --- |
|  | MOI | PHE2 |  | GLA1 |  | PHE2 |  | GLA1 |  |
|  |  | 0.5 | 0.05 | 0.5 | 0.05 | 0.5 | 0.05 | 0.5 | 0.05 |
| Galidesivir | IC<sub>50</sub> | 89.17 | 43.08 | 62.51 | 62.50 | 60.52 | 38.90 | 106.50 | 16.45 |
|  | R<sup>2</sup> | 1.00 | 0.99 | 0.91 | 0.91 | 0.99 | 0.99 | 0.97 | 0.77 |
| Remdesivir | IC<sub>50</sub> | 3.17 | 1.32 | 2.62 | 1.24 | 0.25 | 0.23 | 0.11 | 0.07 |
|  | R<sup>2</sup> | 0.99 | 0.99 | 0.71 | 0.94 | 0.99 | 0.94 | 0.99 | 0.93 |
| HCQ | IC<sub>50</sub> | 24.11 | 4.02 | 12.53 | 3.32 | 92.80 | 42.27 | 13.38 | 9.24 |
|  | R<sup>2</sup> | 0.99 | 0.99 | 0.83 | 0.73 | 0.99 | 0.99 | 0.94 | 0.79 |

## 4 Discussion

Drug repurposing is often attempted in preference to the development of new antiviral compounds as the timeline for initiating trials, safety risks and costs associated with research and development are reduced. They have consequently been a popular choice for clinical studies in COVID-19 since the start of the pandemic. However, despite the demonstration of antiviral activity in vitro, to date, results from clinical trials have been inconclusive or disappointing. Many of these trials were not randomised, were open-label and samples size were often small (<100 patients). The exceptions have been national and international studies such as the RECOVERY^47^ and Solidarity trials^17^ that have provided conclusive data on both antiviral and immunomodulatory therapies. Most studies have been conducted in hospital settings, particularly in intensive care units, examining the drugs’ effects in moderate to severe COVID-19 cases late on in the time course of infection, rather than during early infection. Hospitalised patients commonly present with severe late-stage disease when immunomodulatory treatments are required to reduce hyperinflammation and prevent consequent lung damage. Both dexamethasone and the IL-6 inhibitor genetic variants or virus isolates have been shown to reduce mortality in hospitalised patients^48^. However repurposed antiviral compounds to date have not p rooved as successful.

Prior to clinical trials, the efficacy of candidate therapies is ascertained by antiviral activity in vitro. Here, 29 FDA-approved nucleoside analogues, protease inhibitors, influenza antivirals and other compounds were investigated for anti-SARS-CoV-2 activity in vitro, of which three demonstrated significant antiviral activity. Both the EC_50_ (the concentration of a drug that gives half-maximal response) and the IC_50_ (concentration of an inhibitor where the response is reduced by half) have been widely reported for these compounds in the literature, as indicators of potency, but do vary in their values. These values depend on the conditions of the experiment, such as MOI, cell type and lineage as well as incubation time. On review of in vitro and clinical studies, antiviral activity in cells was not found to necessarily translate to benefit in patients. For example, while HCQ and lopinavir/ritonavir showed strong SARS-CoV-2 activity in cells, they failed to lower mortality rates in hospitalised COVID-19 patients in trials and their use has been discontinued for this patient group^16-18^. Furthermore, some contraindications will require a detailed patient history and continued monitoring, which in milder cases may not be possible^49^. For example, some drug candidates cause a prolonged QT interval and/or arrhythmias, or liver and kidney impairment. It is therefore crucial that such studies are carried out in robust blinded clinical trial settings to fully assess the impact of a given drug on COVID-19 outcomes. A good example of this is HCQ which showed promise in early small-scale trials but was found to cause more harm than benefit, as well as no effect on COVID-19 outcomes^17,18^ by large-scale trials. This may be related to difficulty in achieving adequate doses in vivo.

The anti-viral remdesivir have been shown to improve patient outcomes in one study, although evidence on the latter remains conflicting and no study has shown a significant effect on mortality^50^. Remdesivir (GS-5734) is an adenosine nucleotide prodrug and a broad-spectrum antiviral developed by Gilead, originally used for treatment of Ebola and Marburg virus infections. As an intravenous drug that interferes with viral RNA-dependent RNA polymerase, remdesivir is most effective during early stages of infection, like most antivirals ^51,52^. In 2014, Gilead developed an interest in SARS and MERS, but no clinical trials were carried out due to inadequate patient numbers. Nevertheless, remdesivir was shown to be effective against coronaviruses in vitro^53,54, 26,31 32^ and in vivo^51,55^. Pruijssers et al^33^ showed that remdesivir was effective in mice infected with chimeric SARS-CoV encoding the RNA-dependent RNA polymerase of SARS-CoV-2, while a Gilead study^52^ reported its effectiveness in SARS-CoV-2 infected rhesus macaques. By the end of January 2021, there were 65 trials investigating remdesivir in COVID-19 patients registered in the WHO ICTRP database. A large, randomised trial (n=1063) funded by the NIH showed that remdesivir accelerated recovery in patients with severe COVID-19, reducing median recovery time by four days and improving survival (<u>NCT04280705</u>, ^19^). However, preliminary results from the WHO’s Solidarity trial found no effect on mortality or hospital stay^56^. Similarly, a smaller randomised, double-blind, placebo-controlled multicentre trial study in China (n=237) found no benefits associated with remdesivir and reported that treatment was stopped early because of adverse events in 18 (12%) patients (<u>NCT04257656</u>)^57^.

Like remdesivir, galidesivir (BCX4430, BioCryst) is a novel synthetic adenosine analogue developed for the treatment of Ebola virus and Marburg virus disease. Galidesivir is a broad-spectrum antiviral given intravenously and has activity against several other viruses, including bunyaviruses, arenaviruses, paramyxoviruses, coronaviruses and flaviviruses.^42^ Phase I studies found that it was generally safe and well-tolerated in healthy volunteers. Following promising results against coronaviruses in HeLa cells (MERS-CoV EC_50_=68·5 μM and SARS-CoV EC_50_=57·7 μM)^42^ and a molecular docking study showing that galidesivir can bind tightly the RdRp of SARS-CoV-2^58^, a randomised, double-blind, placebo-controlled clinical trial has commenced (<u>NCT03891420)</u> in Brazil to consider galidesivir for treatment in COVID-19 and yellow fever. Recently, one study reported EC_50_ values under 100 μM for galidesivir, but the experiments used cell culture-adapted SARS-CoV-2_43_. Here, we report IC_50_ values for galidesivir against two isolates of SARS-CoV-2 demonstrating an IC_50_ of between 16·45 to 106.5μM, depending on the viral isolate, MOI and cell line, which is within the same activity range previous demonstrated for SARS-CoV and MERS-CoV.

The anti-influenza drug favipiravir (T-705) is also being investigated for treatment of SARS-CoV-2. Favipiravir is a pyrazinecarboxamide derivative and a pro drug that is converted to a nucleoside analogue within the cell of interest, followed by a series of phosphorylations to produce the active ribofuranosyltriphosphate T-705RTP, which selectively inhibits viral RNA-dependent RNA polymerases^59^. Following a series of clinical trials, favipiravir has been licenced for severe influenza in Japan. While favipiravir has not been tested against SARS-CoV-1 and MERS-CoV, several publications have reported anti-SARS-CoV-2 in vitro, with EC_50_ values ranging from 61·9□M^26^ to 207μM^41^ in VeroE6 cells. In a randomised control trial (n=240), 7-day clinical recovery was more frequent in patients taking favipiravir (71%) compared to umifenovir^60^. Further trials are ongoing to evaluate the efficacy and safety of favipiravir in the treatment of COVID-19, including the Glasgow Early treatment Arm, Favipiravir (GETAFIX) in Scotland ^22,61-63^.

## 5 The way forward

There is an urgent need for the development of novel antivirals to treat COVID-19 and tackle the pandemic. Since antiviral drugs are most likely to be effective early in the disease process, what are ideally required are antiviral therapies that can be taken orally and are well enough tolerated to be widely available for use in the community very soon after onset of symptoms. Currently the drug which has closest to this product profile is Favipiravir, which has had success in early clinical trials ^60^ but had no antiviral activity in our hands in vitro. A bespoke small molecule with a lower EC_50_ than Favipiravir might have a greater chance of success.

An alternative direction for the development of antiviral therapeutics is anti-SARS-CoV2 monoclonal antibodies (mAbs). For example, the S protein blockers casirivimab/imdevimab, also known as REGN10933/REGN10987^64^ and Bamlanivimab (LY-CoV555)^65^ which have both been issued with emergency use authorisation by the FDA, for use in non-hospitalised patients not requiring oxygen. Also under investigation in animal models, is CT-P59, which targets the receptor binding protein of SARS-CoV2^66^. Casirivimab/imdevimab showed significant reductions in virus levels and were associated with significantly fewer medical visits^64^. Bamlanivimab in combination with etesevimab decreased SARS-CoV2 log viral load by day 11 (BLAZE-1 study) ^65^.

The EC_50_s for mAbs are promising (e.g. 31pM for casirivimab/imdevimab^67^, however, mAb preparations have some disadvantages that will limit their use. They have to be administered intravenously, and adverse events can include serious hypersensitivity reactions and anaphylaxis. These attributes preclude the possibility of using them as community-based therapies. Further, monoclonal antibodies tend to select for escape viral variants, which may limit their useful life span. Using monoclonal antibodies in combination is an attempt to limit escape viral variants. For example, escape variants have been documented when using either casirivimab or imdevimab individually, but so far not when used in combination ^67^.

Repurposed compounds have been evaluated in hundreds of studies with limited clinical success, there remains an urgent need both for the development of novel antivirals to tackle severe disease and for a product that is safe enough and logistically practical for use in the critical early days of the disease process. It is time to think beyond the immediate emergency and fund the development of bespoke drugs for long-term solutions.

## 6 Acknowledgements

We would like to acknowledge the kind gift of cell lines from Prof. Michèle Bouloy (Institut Pasteur, France) and Prof. Rick Randall (University of St Andrews, UK). In addition, we would like to acknowledge the support of members of the Centre for Virus Research who provided reagents to enable the work within this study, Suzannah Rihn and Matthew Turnbull for the development of the various cell lines used, and Arthur Wickenhagen for the SARS-CoV-2 virus stocks.

## Funding

AK: UK Medical Research Council funding (MC_UU_12014/8).

DK: UK Medical Research Council MRC Precision Medicine Training Grant (MR/N013166/1-LGH/MS/MED2525).

MES and CD: Medical Research Council core award (MC_UU_1201412)

## Declaration of interests

The funders had no role in study design, data collection and analysis, decision to publish, or preparation of the manuscript. JTS is a chief investigator on the GETAFIX trial (ISRCTN31062548), other authors have declared no competing interests.

## Supplementary Material

**Supplementary Table 1.**
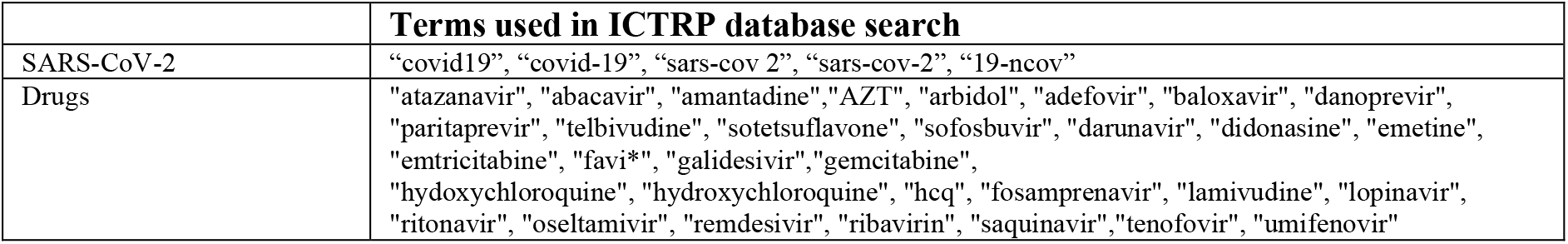
Terms used for searching the ICTRP database. Note that hydroxychloroquine (HCQ) and favipiravir were often misspelled, hence the more flexible terms.

**Supplementary Table 2.**
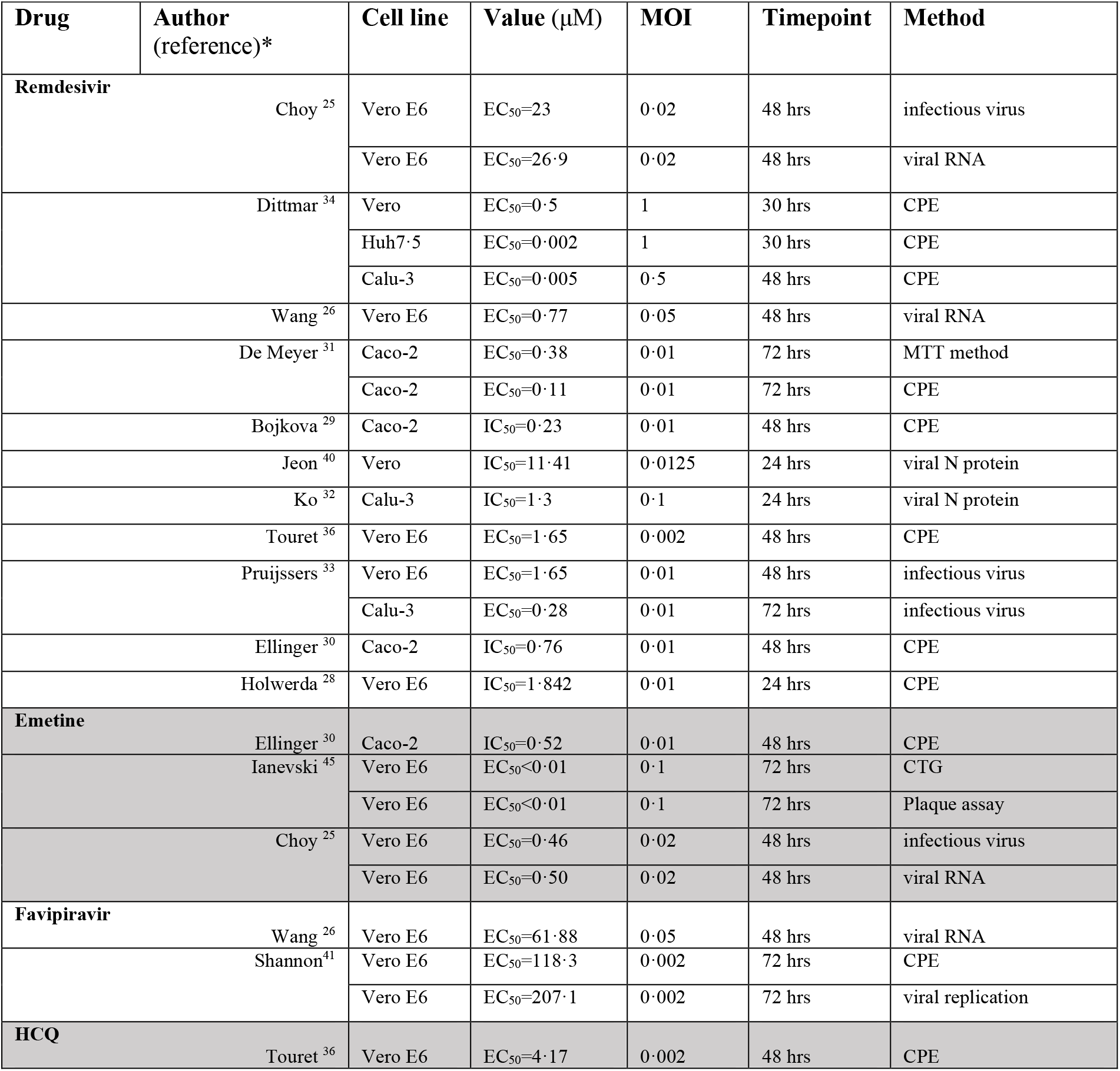
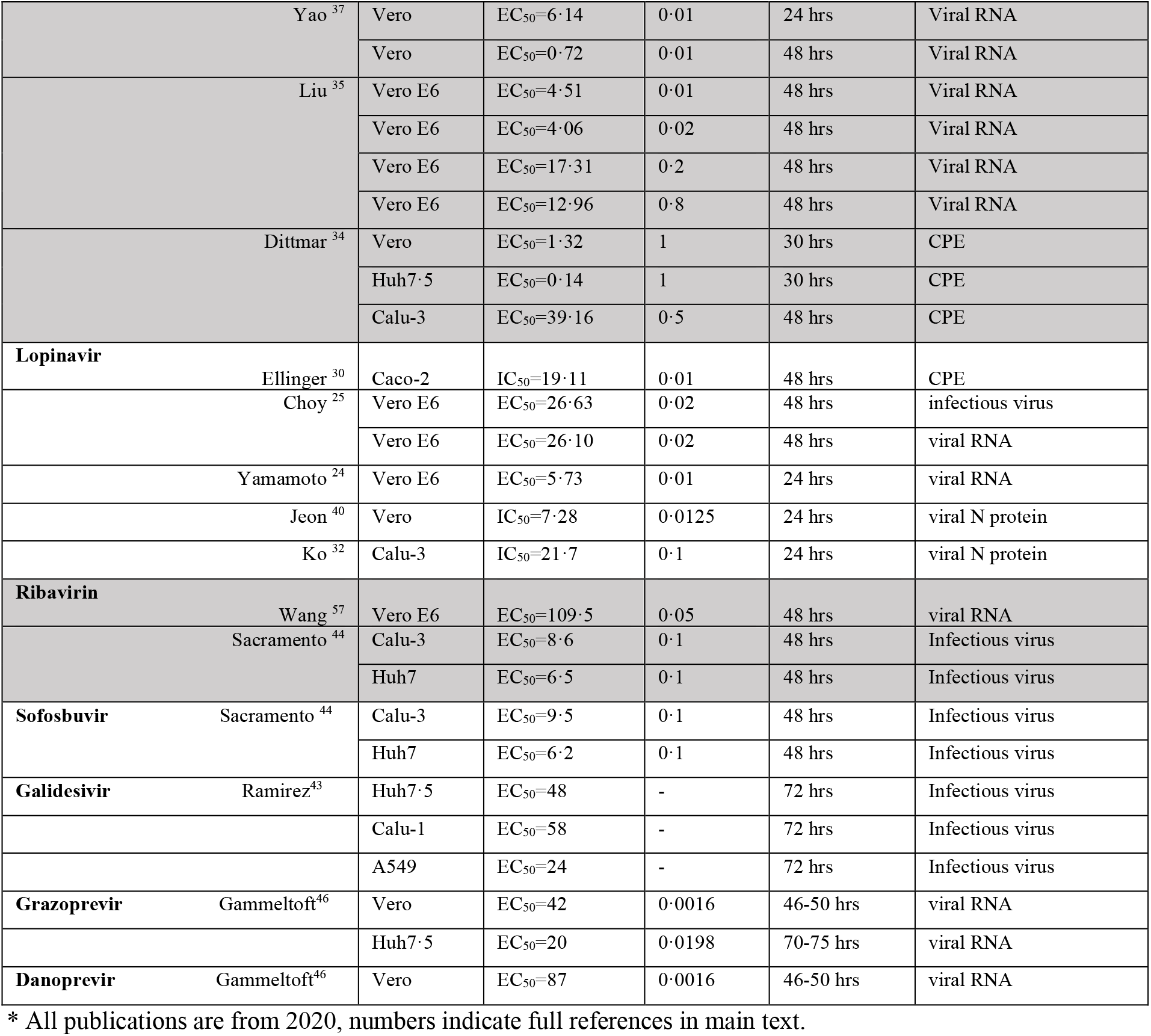
Source and details of EC_50_/IC_50_ values used for Figure 3, including multiplicity of infection (MOI). Table adapted from Stanford University’s Coronavirus Anti-Viral Research Database (https://covdb.stanford.edu). CPE = cytopathic effect.

**Supplementary Table 3.**
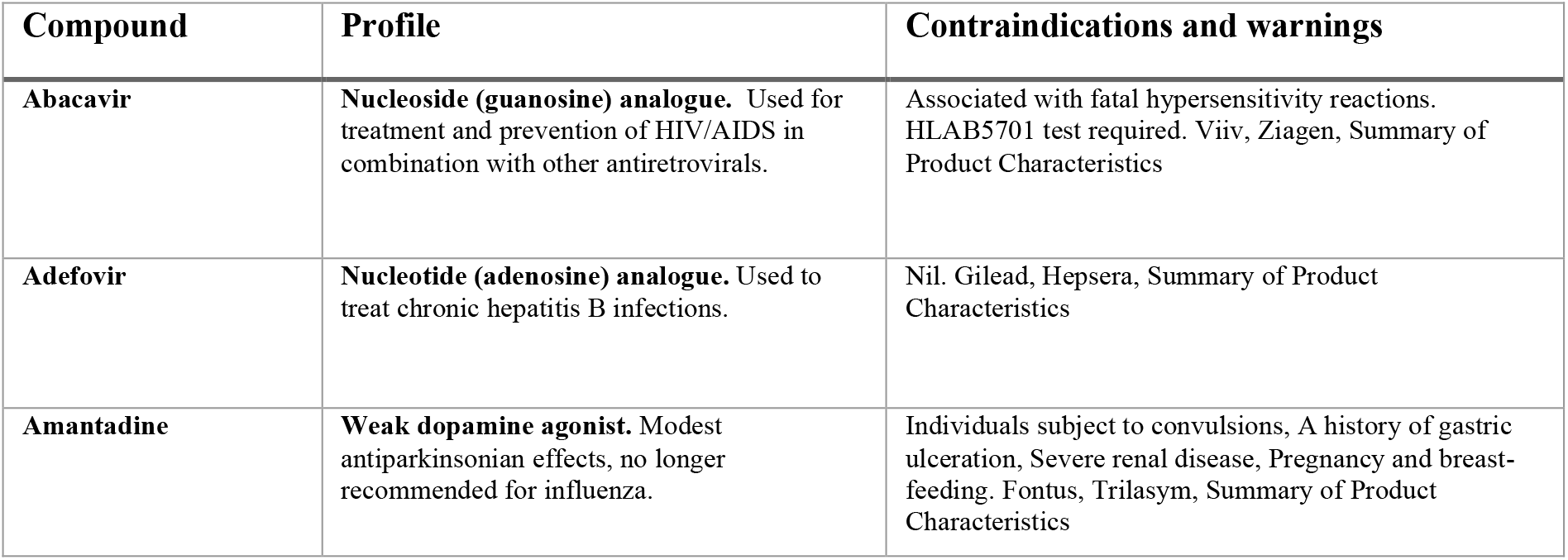
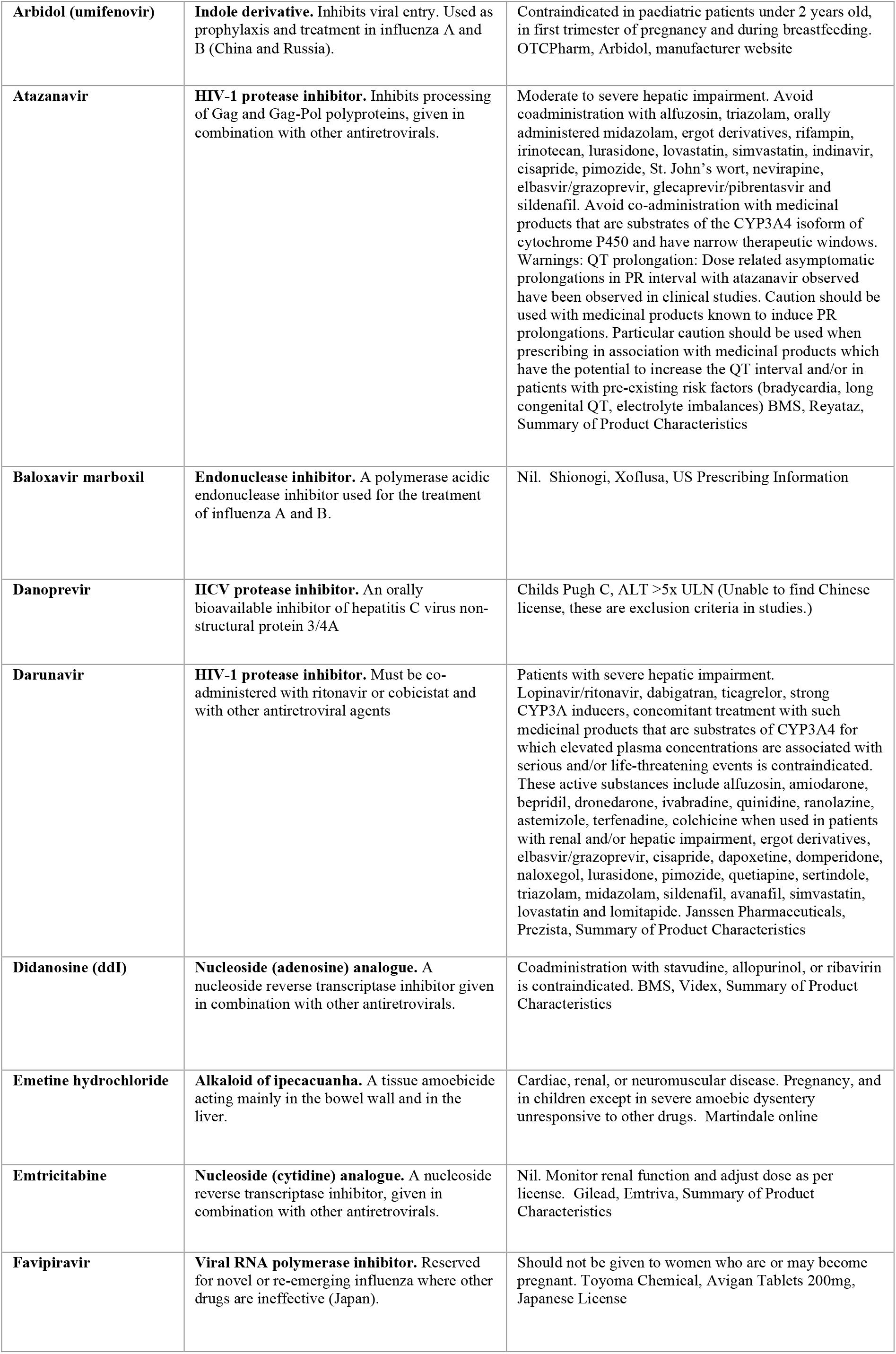
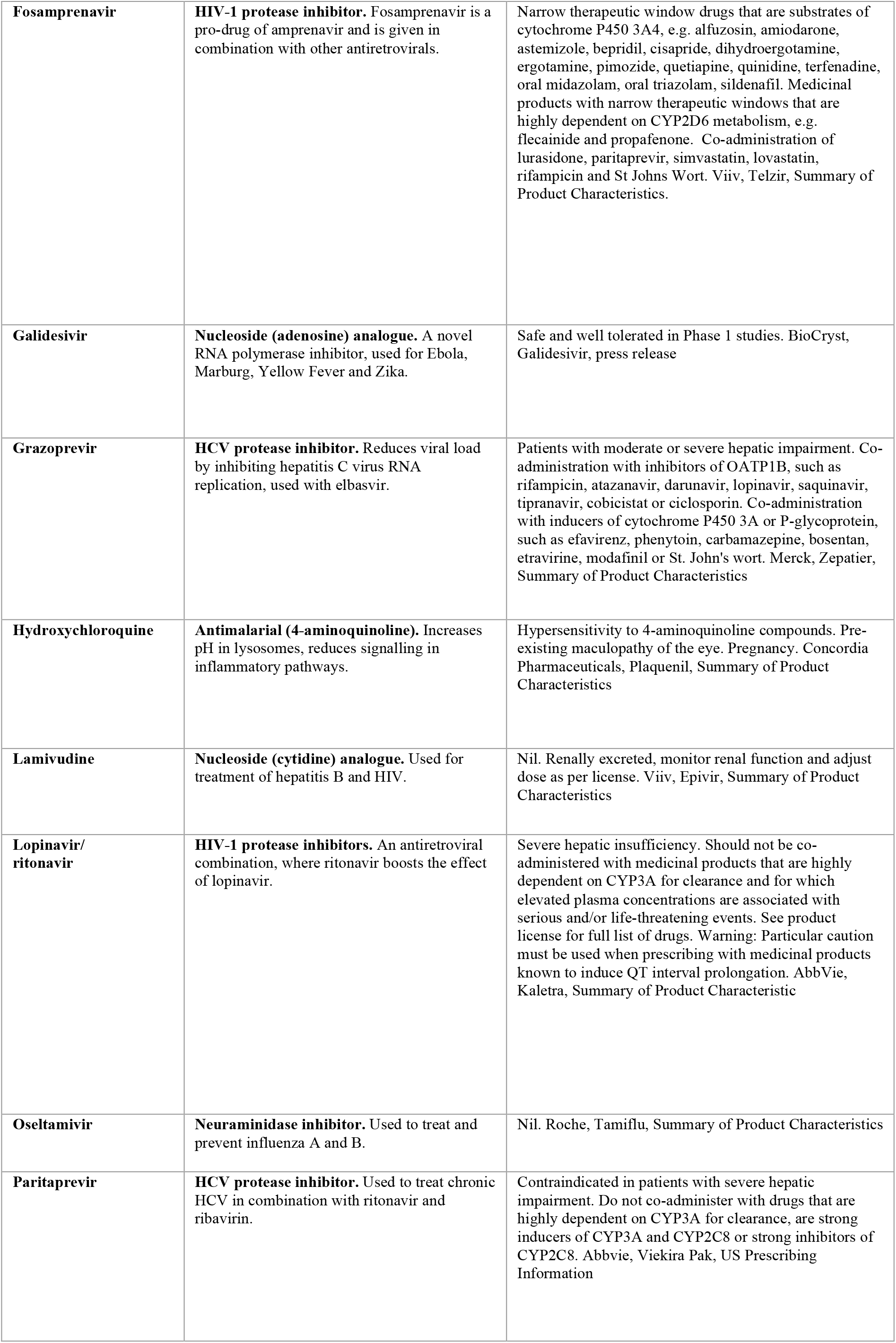
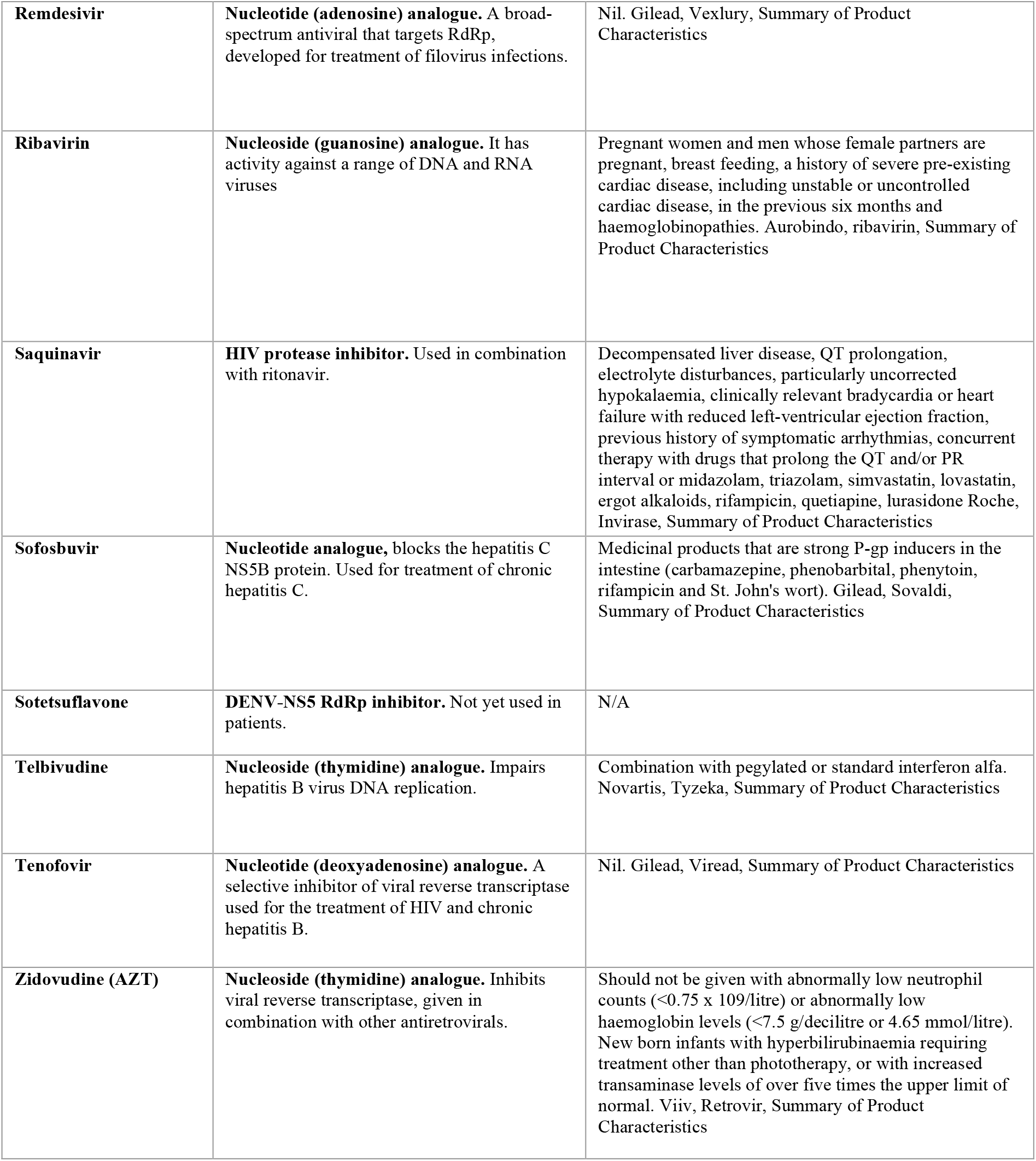
Contraindications and warnings associated with 29 FDA-approved repurposed compounds.

**Supplementary Figure 1.**
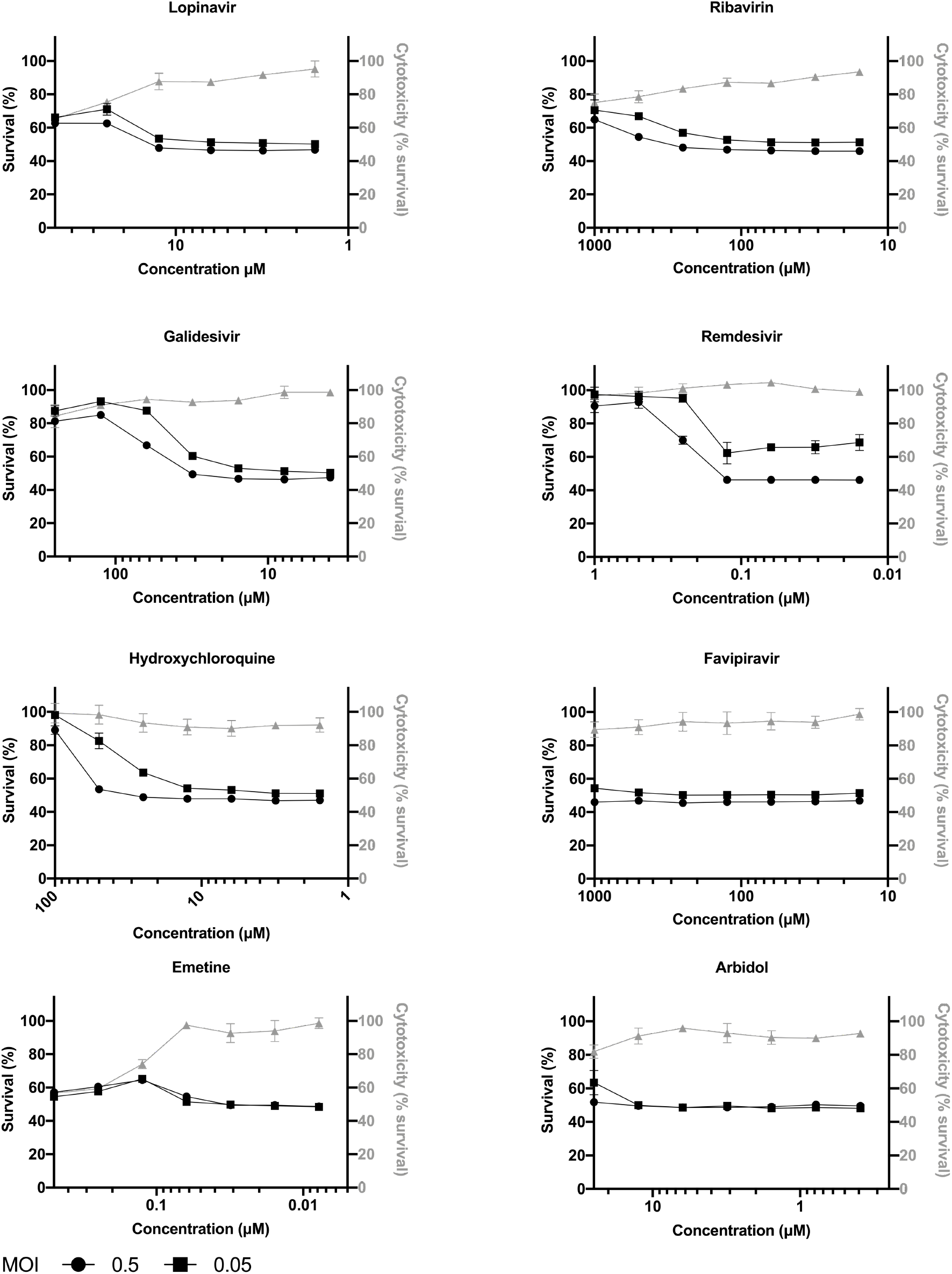
Assessment of SARS-CoV-2 inhibition by active FDA approved antiviral compounds. A549 cells, overexpressing Ace2 and BVDV Npro protein, were treated for three hours with varying concentrations of antiviral compounds before the addition of SARS-CoV-2 PHE2 isolate at an MOI of 0·05 and 0·5. The cells were fixed with 8% formaldehyde 72 hours post-infection and stained with Coomassie blue. Percentage cell survival was determined by comparing the intensity of the staining to uninfected wells. The data shown are representative figures of 3 independent repeats. The mean percentage cell survival of the DMSO control infected with an MOI of 0·05 and 0·5 was 51.08% (SD 1·117) and 47·35% (SD 0·764), respectively. The cytotoxicity of the compounds is depicted by the grey line and were determined by comparing the intensity of staining to untreated wells.

**Supplementary Figure 2:**
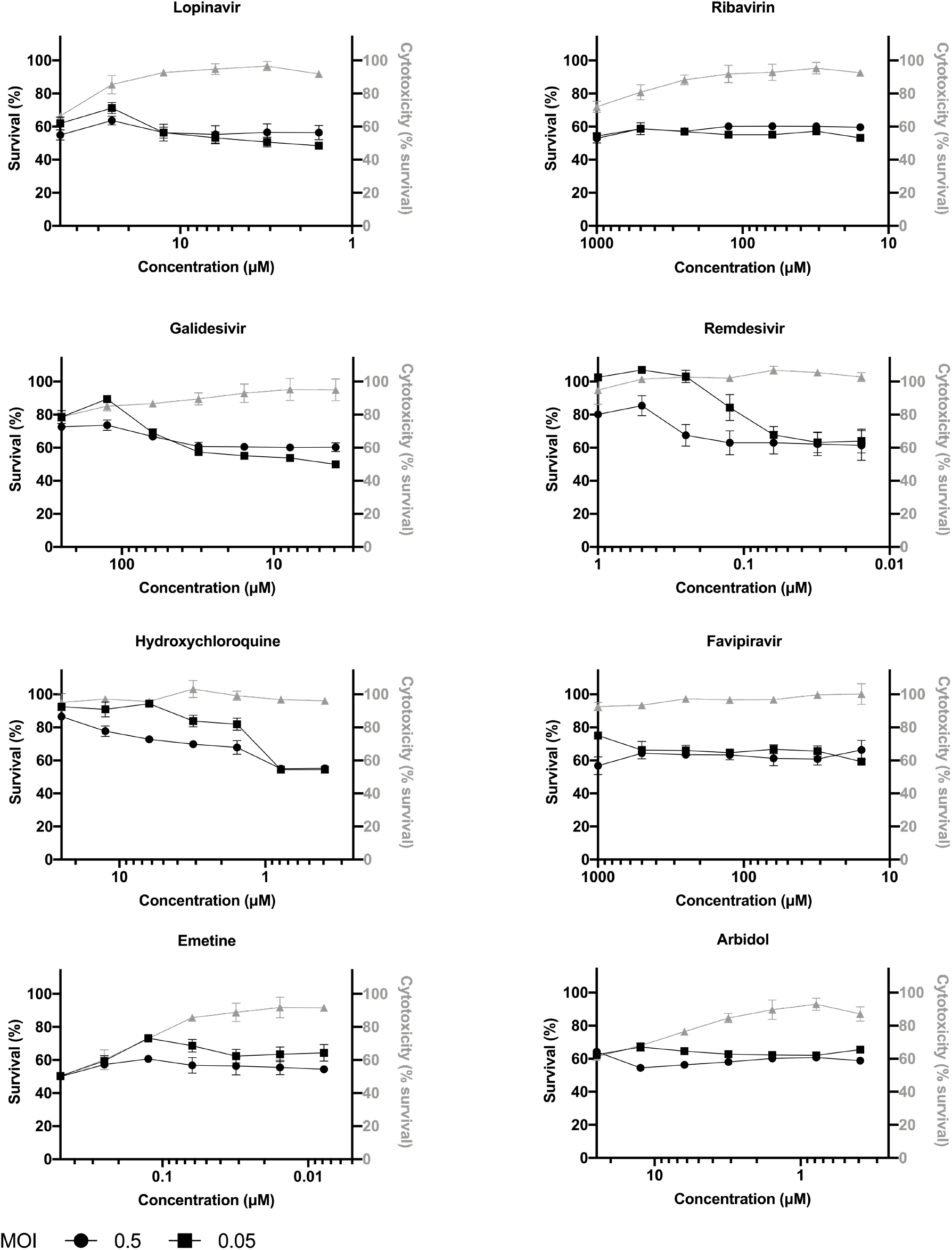
Assessment of SARS-CoV-2 inhibition by active FDA approved antiviral compounds. VeroE6 cells, overexpressing Ace2, were treated for three hours with varying concentrations of antiviral compounds before the addition of SARS-CoV-2 GLA1 isolate at an MOI of 0·05 and 0·5. The cells were fixed with 8% formaldehyde 72 hours post-infection and stained with Coomassie blue. Percentage cell survival was determined by comparing the intensity of the staining to uninfected wells. The data shown are representative figures of 3 independent repeats. The mean percentage cell survival of the DMSO control infected with an MOI of 0·05 and 0·5 was 50% (SD 0·502) and 43·45% (SD 0·642), respectively. The cytotoxicity of the compounds is depicted by the grey line and were determined by comparing the intensity of staining to untreated wells

**Supplementary Figure 3.**
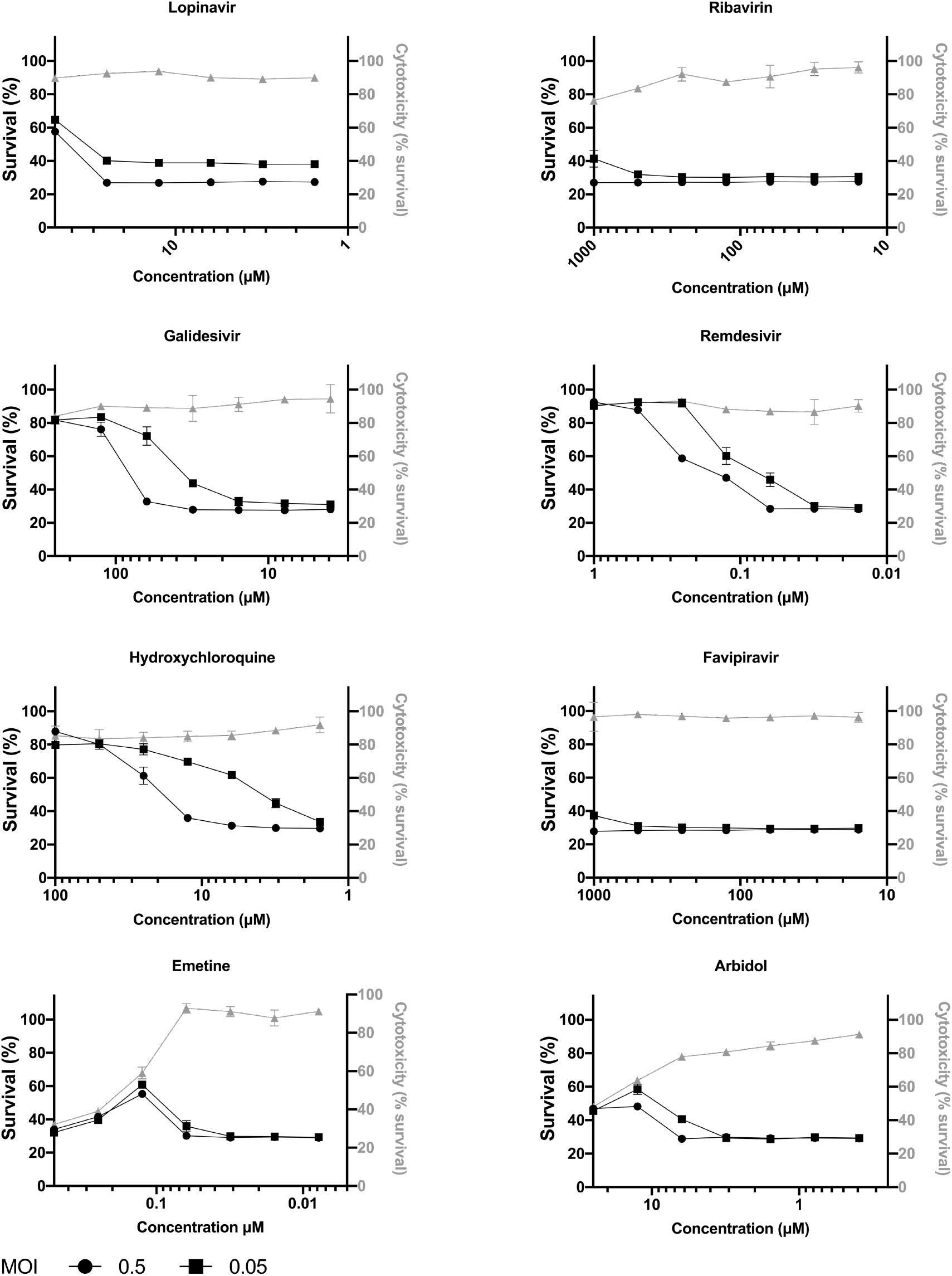
Assessment of SARS-CoV-2 inhibition by active FDA approved antiviral compounds. VeroE6 cells, overexpressing Ace2, were treated for three hours with varying concentrations of antiviral compounds before the addition of SARS-CoV-2 PHE2 isolate at an MOI of 0·05 and 0·5. The cells were fixed with 8% formaldehyde 72 hours post-infection and stained with Coomassie blue. Percentage cell survival was determined by comparing the intensity of the staining to uninfected wells. The data shown are representative figures of 3 independent repeats. The mean percentage cell survival of the DMSO control infected with an MOI of 0·05 and 0·5 was 28·83% (SD 0·673) and 27.52% (SD 0·193), respectively. The cytotoxicity of the compounds is depicted by the grey line and were determined by comparing the intensity of staining to untreated wells

**Supplementary Figure 4.**
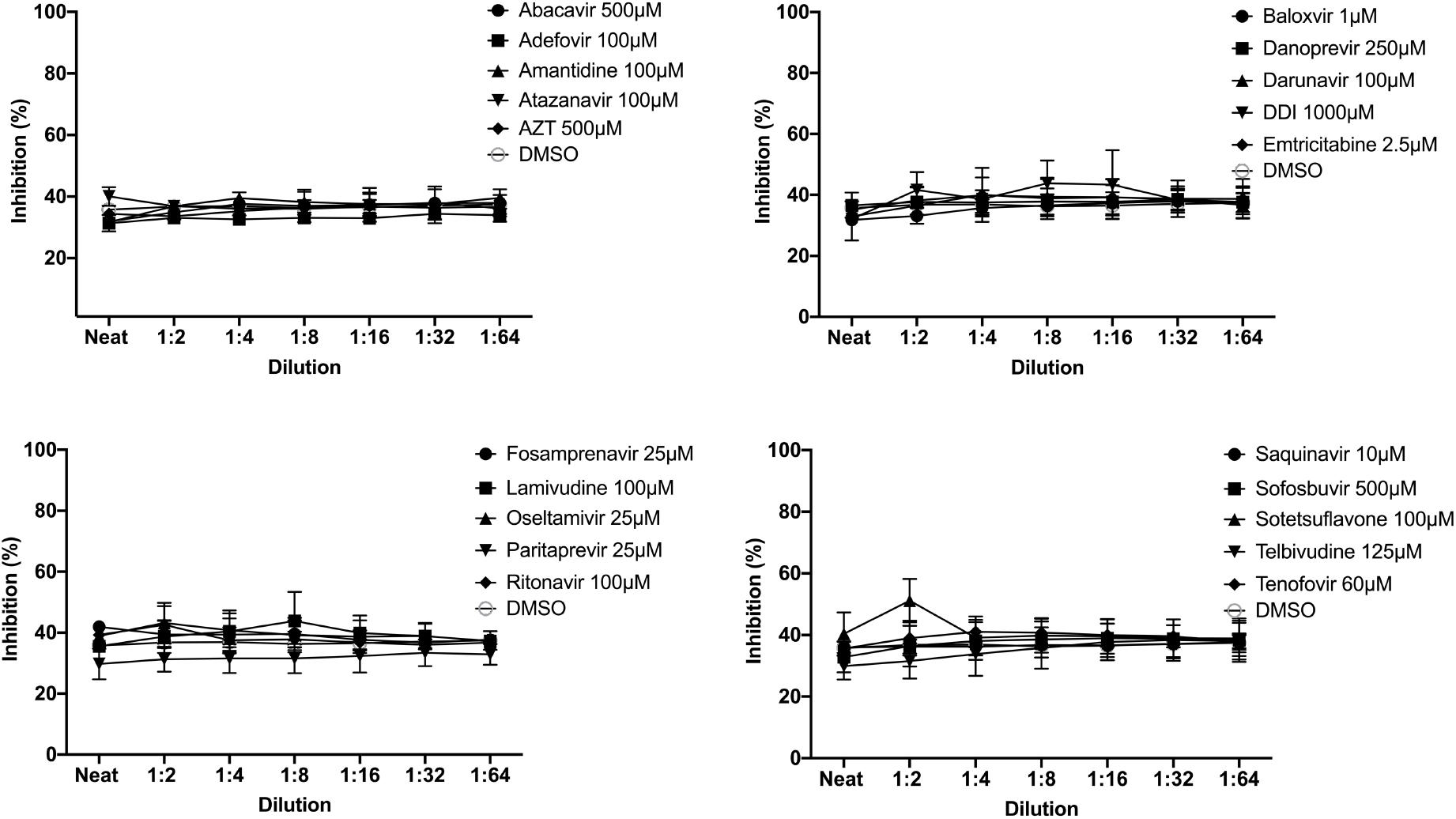
Assessment of SARS-CoV-2 inhibition by active FDA approved antiviral compounds. A549 cells, overexpressing Ace2 and BVDV Npro protein, were treated for three hours with varying concentrations of antiviral compounds before the addition of SARS-CoV-2 PHE2 isolate at an MOI of 0*·*5. The cells were fixed with 8% formaldehyde 72 hours post-infection and stained with Coomassie blue. Percentage cell survival was determined by comparing the intensity of the staining to uninfected wells. The data shown are representative figures of 2 independent repeats.

